# Modelling Metacognition: A Joint Prediction-Confidence Model for Predictive Inference Task Data

**DOI:** 10.64898/2026.08.14.744790

**Authors:** Esteban Félez Martínez, Peter Thestrup Waade, Jakob Heinzle, Alexander J. Hess

## Abstract

Metacognition is the ability to reflect on and evaluate our own cognitive processes. It is often altered in psychopathology. Yet, the computational mechanisms underlying these alterations remain unclear. In this work, we extend Hierarchical Gaussian Filter (HGF) models to jointly fit trial-by-trial predictions and confidence ratings in a predictive inference task, providing an individualised characterisation on metacognitive processing. Applying our cognitive computational model to a large subclinical open dataset (N=430), we are able to achieve, on average, excellent fit of prediction responses 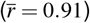 and a moderate to good fit of confidence ratings 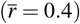. Analysis of experimental change-points revealed that our model accurately captures confidence self-reports dynamics around these change-points. Posterior parameter estimates reveal a negative effect of sensory input prediction errors and a positive effect of sensory input prediction precision on confidence ratings, respectively. In addition, we replicate state-of-the-art findings related to compulsivity as measured by a transdiagnostic factor score, such as inflated confidence and a decoupling of action updates (here, prediction errors) and confidence in compulsivity. These results demonstrate the robustness of our methodology and the potential of joint prediction-confidence modelling to uncover latent metacognitive alterations in psychopathology.

## Introduction

Metacognition is the ability to reflect on and evaluate our own cognitive processes, playing an important role in different types of communication, reading and oral comprehension, problem solving, social cognition and cognitive behaviour modification^1^. A common way to assess metacognition in humans is through confidence self-judgments, which are intended to capture the subjective feeling of being correct about a decision or statement^2,3^. Computational cognitive models represent one way of formalising concrete mechanistic hypotheses depicting actual mental processes that are empirically testable. They offer a framework to investigate human behaviour by constructing such models and applying them to suitable, behavioural data^4–7^. Hence, cognitive computational models of metacognitive belief formation represent a crucial step towards advancing our understanding of metacognition.

Abnormalities in confidence judgments are ubiquitous in the subclinical stages of psychiatric disorders and scale positively with symptom severity^3^. In the field of computational psychiatry, computational cognitive models of mental processes are used to quantify (aberrant) belief formation in psychopathology. Concretely, such models might help uncover metacognitive deficits present in conditions such as Obsessive Compulsive Disorder (OCD); observed both during memory tasks^8–10^ and predictive inference tasks^11,12^, where compulsivity was shown to be associated with inflated confidence ratings and a decoupling between task performance and confidence self-reports. Developing a computational cognitive model of metacognition of learning can enable the quantification of both normative variability in learning and confidence formation, as well as the identification of deviations related to clinical profiles.

Existing theoretical accounts of metacognition conceptualise the experience of confidence as reflecting the Bayesian posterior probability of a choice being correct^13–16^. In parallel, numerous models have been proposed to mechanistically describe confidence in population samples, most of them based on signal-detection theory^17–19^or sequential-sampling models such as the drift-diffusion model^20–22^. However, these accounts mainly focus on metacogntion of perception, memory, or decision-making and many of these studies focus on population-averaged behaviour and often do not examine trial-by-trial variance in confidence ratings. An exception marks the work using data from a predictive inference task^11,12^, which focuses on metacognition of probabilistic learning. In this work, the authors computed the behaviour of a quasi-optimal Bayesian learner based on the sequence of observations experienced by each participant. The resulting parameters of this ideal observer model (e.g., model-based prediction error or change-point probability) were regressed against participants’ prediction responses and associated confidence ratings. Although this approach allowed for a comparison of statistically optimal and real human behaviour, ideal observer models cannot capture inter-individual variability in belief updating and how these give rise to behaviour, including metacognitive self-judgements. Hence, an individualised, trial-by-trial account of belief updating, responses and confidence ratings in this task may help uncover the different mechanisms and biases underlying learning metacognition.

Hierarchical Gaussian Filtering (HGF)^23–25^is a widely used form of computational cognitive models relying on hierarchical Bayesian belief updating to describe perception and learning. As all cognitive models, applying them to behavioural data allows for a mechanistic characterisation of inter-individual variability in inference and learning, which can then be related, for example, to clinical markers of psychiatric conditions (for applications in computational psychiatry, see e.g. Refs^26–34^). Bayesian methods are usually used to fit this cognitive model to behavioural data, in what has been referred to as the Bayesian ‘observing the observer’ framework^5,35,36^. If HGF models were extended to describe the relationship between hierarchical Bayesian belief updating and confidence formation, this could enable valuable insights into metacognitive processes related to learning and perception. The simplest form of such an extension requires the specification of an appropriate response or action model providing a mapping from inferred beliefs of an agent to observed responses recorded during an experiment. Specifically, multivariate action models represent one possible avenue of implementation for the application of HGF models to learning and metacognition. Hess et al. recently introduced multivariate action models allowing joint fitting of binary choice data alongside participants’ reaction times in an associative learning task^37^. Multivariate action models possess several advantages over classical univariate action models often used in HGF applications: On the one hand, they provide a more holistic account of action generation. On the other hand, harnessing information from different response modalities can improve the robustness of statistical inference.

In this work, we introduce a novel joint prediction-confidence model in the HGF framework explaining trial-by-trial prediction responses and confidence ratings in the flying particle predictive inference task^12^, also known as the jumping Gaussian estimation task (JGET). For the perceptual component of our model, we use a previously developed HGF variant for this task, the JGET HGF^26^. We augment the perceptual component with an action model capable of simulating and fitting prediction responses and confidence ratings jointly. This enables the quantification of individual-specific metacognitive belief formation in the context of learning. We validate our model using both simulations and model inversion on empirical data from Ref.^12^ (N=430). In addition, we demonstrate the utility of our approach for computational psychiatry by relating our model-based estimates of learning and confidence formation during the task to questionnaire-based reports of psychiatric symptoms.

## Methods

### Dataset

For our analysis, we used a publicly available dataset from Ref.^12^ (https://osf.io/2z6tw/). Here, we briefly summarise the aspects of the dataset relevant for our application (see the original publication for a more detailed description). The dataset contains behavioural data from 437 participants who performed a predictive inference task consisting of 300 trials adapted to web-based testing (Fig. 1a). After the behavioural task, participants completed an IQ-test and a range of self-report psychological questionnaires (209 questions in total, described in detail in Ref.^12^). Questionnaire data were mapped onto three transdiagnostic factor scores following the approach presented in Refs.^12,38^, yielding scores for Compulsive and Intrusive Thought (CIT), Social Withdrawal (SW) and Anxious Depression (AD). At the time of task completion, 41 participants (9.38%) were being medicated due to mental health issues. For our analysis, we excluded participants who showed consistently low confidence ratings with very little variation (defined as having a median below 5 and a standard deviation below 10), leaving a total of 430 participants, as these participants may not have meaningfully engaged with the task, may have misunderstood the rating scale, or may have responded in a non-differentiated manner (e.g., repeatedly reporting the same confidence). Fig. 1b contains histograms of transdiagnostic factor scores (CIT, AD, SW), demographic variables (age, gender), and average self-reported confidence ratings per participant 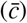 in the dataset. Examples of predictive inference task data for a given participant are shown in Fig. 1c.

**Figure 1.**
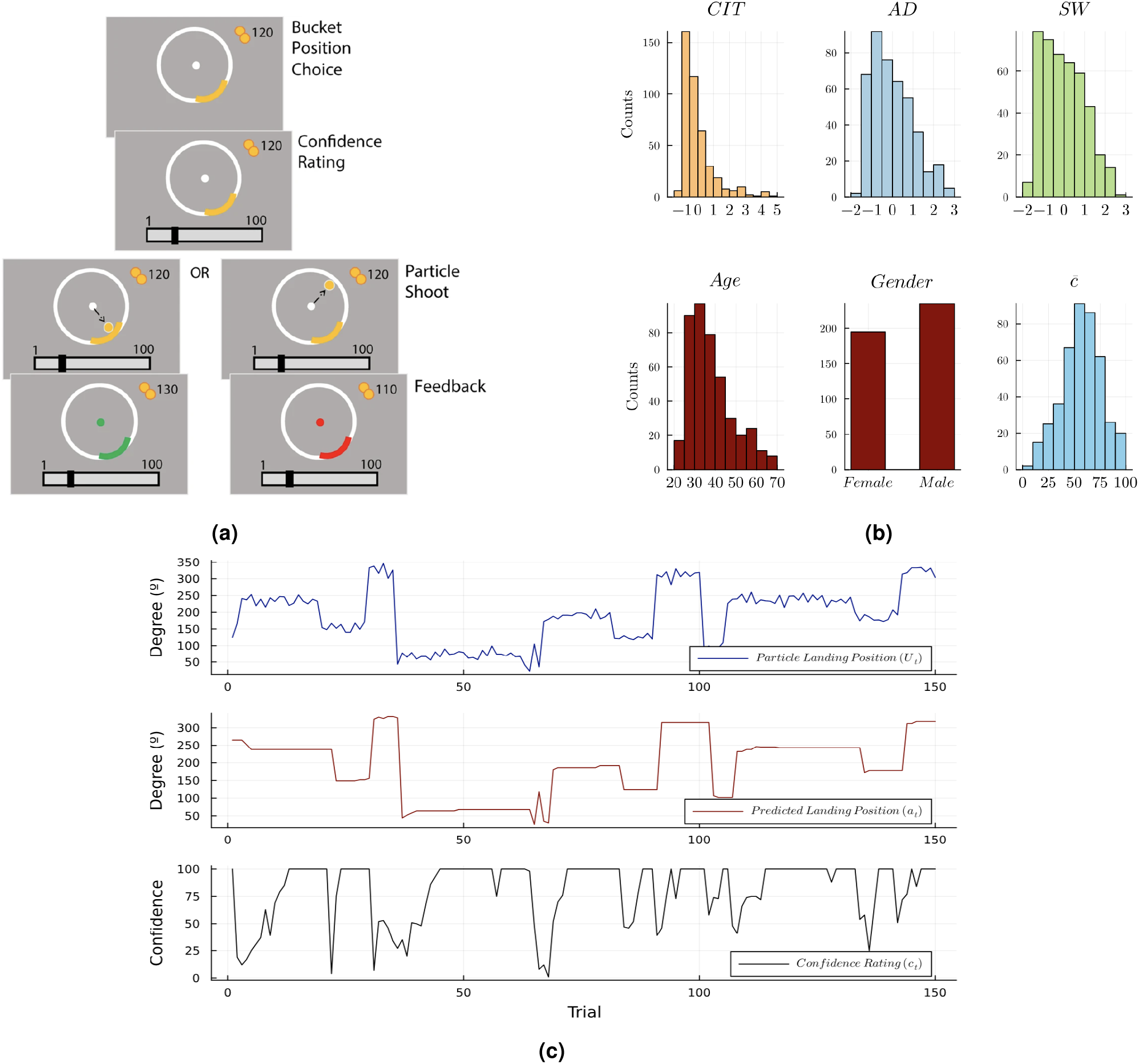
(a) Visual summary of a single trial in the predictive inference task. Figure reproduced from Ref.^12^ under a CC-BY license. (b) Histograms showing the distribution of transdiagnostic factor scores (Compulsive and Intrusive Thought; CIT; Social Withdrawal, SW; and Anxious Depression, AD) and demographic variables (age, gender) as well as average confidence ratings across 150 trials 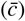 for N=430 participants included in our analysis. The transdiagnostic factor scores were z-scored to be consistent with Ref.^12^. (c) Example time series of particle landing positions (*U*_*t*_), predicted landing positions (*a*_*t*_), and confidence ratings (*c*_*t*_) for participant 53.

#### The predictive inference task

First introduced by Vaghi et al.^11^, the flying particle predictive inference task has been used to study learning and belief updating under uncertainty^11,12,26^. In this task, participants were required to predict the future landing position of a particle in a circular space and rate their relative confidence in these predictions. A visual summary of a single trial *t* in the predictive inference task is shown in Fig. 1a. At the start of each trial, participants were instructed to place a bucket on the perimeter of a circle to catch a particle flying from the centre to the edge of the circle. Once the location of the bucket was set (predicted landing position, *a*_*t*_), participants had to report their confidence (*c*_*t*_, on a scale of 1 to 100) that the particle would land in the bucket. After this, participants received visual feedback about the particle’s actual landing position (*U*_*t*_), and hence also about correctness of their prediction. On each trial, the landing position of the particle was sampled from a Gaussian distribution with a fixed standard deviation of 12 degrees and a mean which changed abruptly several times throughout the experiment. In the task, there were two hazard rate conditions defining the probability of a change in the mean of the observation generating distribution (change-point) over a stretch of 150 trials each: stable (hazard rate = 0.025, 4 change-points) and volatile (hazard rate = 0.125, 19 change-points). For simplicity, we only used data from the volatile condition in our analysis.

#### Preprocessing of circular data

As described above, both inputs (particle landing positions *U*) and predictions (bucket positions *a*) were circular variables that cover the circular range of 0º–360º. To account for the circular nature of the task data in our modelling, we unrolled the landing positions and predictions to put them on the real line (see Supplementary Fig. S8 for a depiction of the algorithm used for this). After unrolling, we standardised (z-scored, i.e., mean subtracted and divided by the standard deviation) both the inputs and predictions. In addition, confidence reports were normalised to the [0,1] interval.

### Computational Cognitive modelling

To ensure transparency and robustness of our model development and analysis pipeline, we followed the general steps of Bayesian workflow for generative modelling outlined in Refs.^37,39^. In other words, we engaged in iterative model development prior to applying the model to empirical data. This included the specification of likelihood and priors, prior predictive checking, the choice of approximate Bayesian inference algorithm and concurrent validation of computation using simulations. After inversion of our model on the empirical dataset, we carefully evaluated our model fits and engaged in posterior predictive checking. In what follows, we start by detailing the cognitive computational model consisting of a perceptual model and a response model^35^. Fig. 2 provides a graphical representation of our computational cognitive model for predictive inference task data.

**Figure 2.**
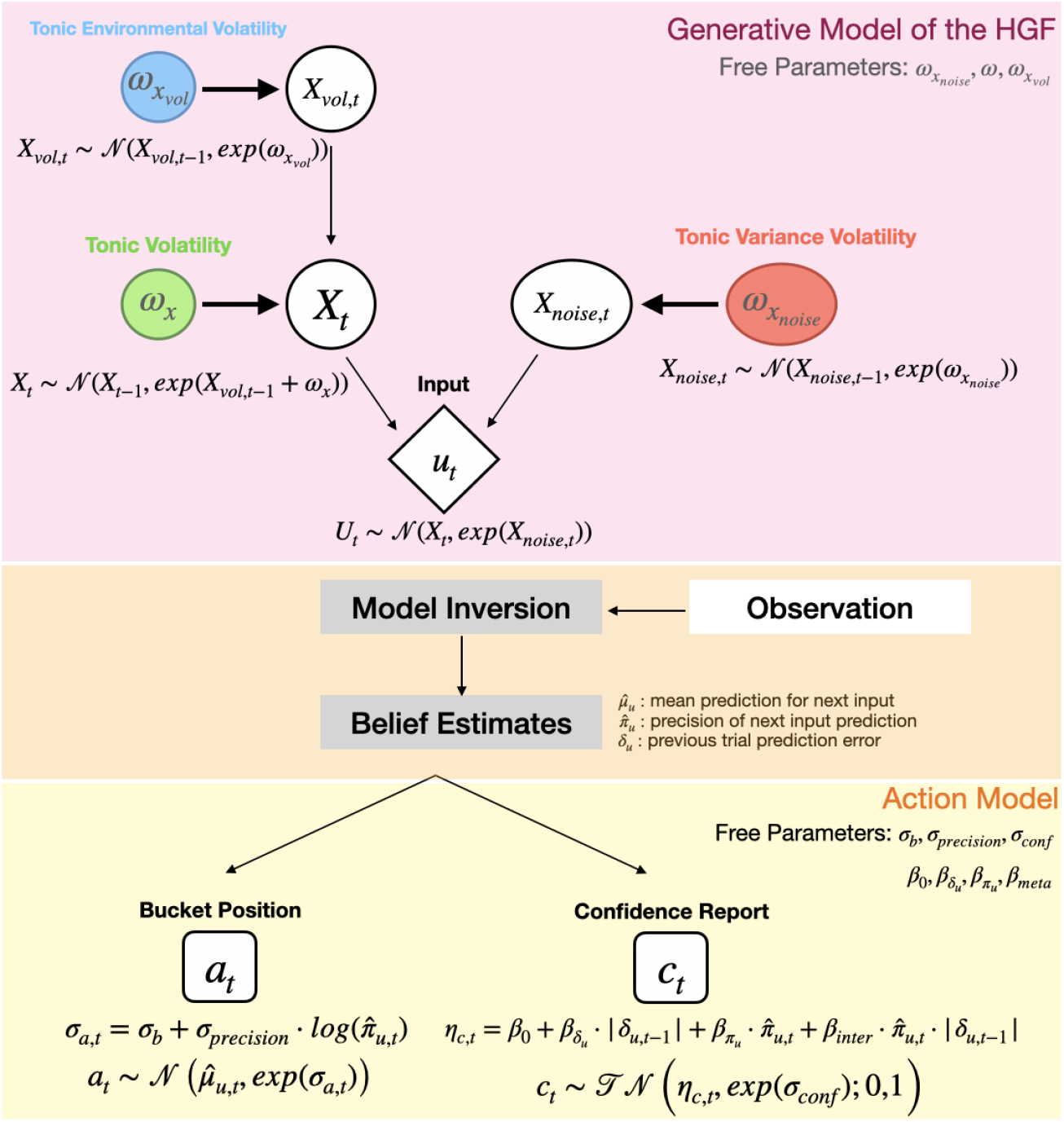
Graphical representation of the perceptual model (the jumping Gaussian estimation task Hierarchical Gaussian Filter; JGET HGF) and the multivariate action model for predictions (*a*) and confidence ratings (*c*). The generative model underlying the JGET HGF comprises three continuous state nodes (*X, X*_*vol*_, *X*_*noise*_) and one continuous observation node (*U* ; particle landing positions). The variational inversion of the generative model yields trial-by-trial update equations for beliefs about the evolution of states over time. The action model uses the perceptual model under its inversion and maps from inferred beliefs to observed behaviour (i.e., bucket positions *a* and confidence ratings *c*).

#### Perceptual model: the Hierarchical Gaussian Filter

We employed Hierarchical Gaussian Filtering (HGF)^23–25^to model the participants’ trial-by-trial belief updating. HGF is a variational Bayesian predictive processing model of perception and learning. The underlying generative model describing the evolution of states in a dynamic environment consists of interlinked Gaussian random walks, a useful assumption when the true dynamics of the environment is not known. Approximating Bayesian inference for this generative model with variational message passing update equations yields single-step precision-weighted prediction error-based updates similar to classical predictive coding architectures^40^. In other words, trial-by-trial update equations resulting from variational inversion of the generative model of the HGF provides a neurobiologically plausible and computationally tractable model of how the mind might perform online Bayesian inference. In its generalised formulation (gHGF^25^), it allows for flexible network construction such that nodes representing separate environmental states can be linked: for example drift, volatility or observation noise can depend on other states that are also changing in time.

For this study, we used the JGET HGF introduced by Mikus et al.^26^ to model belief updating in the predictive inference task. The JGET HGF^26^ simultaneously tracks the position of the target and the observation noise in order to adjust belief updates according to the precision of the observations. In the generative model of the JGET HGF, a continuous input *U*_*t*_, represented by an input node, is assumed to be generated at every trial by a Gaussian distribution with changing mean and variance. The variance of the Gaussian distribution is controlled by a different state, the noise parent *X*_*noise,t*_, represented by a continuous state node. The mean of the input node’s Gaussian distribution depends on a value parent *X*_*t*_, represented by a different continuous state node. *X*_*t*_ evolves according to a Gaussian random walk, where the volatility (i.e., the variance of the Gaussian random walk) is controlled by a tonic noise parameter *ω*_*x*_. In addition, the volatility of *X*_*t*_ is influenced by its volatility parent *X*_*vol,t*_, another continuous state node evolving according to a Gaussian random walk with step size exp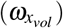. The generative model equations of our model (cf. Fig. 2) are presented together with the update equations in the HGF generative model equations and HGF update equation sections of the Supplementary Materials.

#### Action model: joint prediction response and confidence rating model

In each trial *t*, our action model used the following variables that represent perceptual model belief states to form predictions for the measured behavioural responses (bucket position or prediction *a*_*t*_, confidence rating *c*_*t*_): the mean of the prediction for the next input 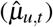, the precision of the next input prediction 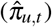, and the previous trial prediction error (*δ*_*u,t−*1_). We modelled the reported prediction *a*_*t*_ to follow a Gaussian with mean 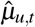 and standard deviation exp(*σ*_*a,t*_):

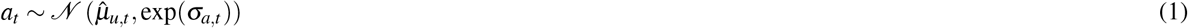

The trial-by-trial noise term *σ*_*a,t*_ depended on a constant baseline noise parameter *σ*_*b*_ and the precision of the input prediction 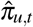. The latter term reflects the influence of certainty on the stochasticity of the bucket position *a*_*t*_, which was weighted by another constant *σ*_*precision*_:

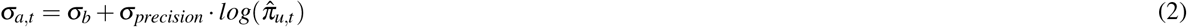

Similarly, we modelled the confidence ratings using a truncated normal distribution with mean *η*_*c,t*_ and standard deviation exp(*σ*_*c*_), on the interval [0,1]:

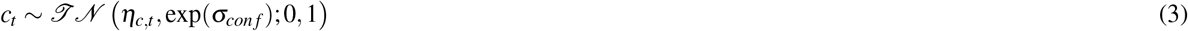

where *η*_*c,t*_ represents a linear combination of different belief states including an intercept *β*_0_, the absolute prediction error of the previous trial |*δ*_*u,t−*1_|, the precision of the input prediction 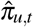, and a linear interaction term between the two:

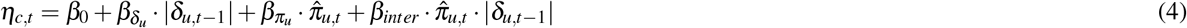

Because empirical confidence ratings were bounded within the (0, 1) interval in the predictive inference task, the latent confidence expectation *η*_*c,t*_ was constrained to the open interval (0, 1), strictly excluding the boundary values of 0 and 1. This was done using a response function which is piecewise linear, with slope 1 in the range (0,1) and slope 0 otherwise. This prevented numerical instability at the boundaries where the probability density function of the truncated normal distribution can diverge or become computationally undefined, solving some sampler convergence issues that were observed in the initial parts of this study. It also avoids degenerate extreme-value states while accurately reflecting the non-infinite precision of human self-evaluations.

In total, the full cognitive computational model (perceptual and action model) contained ten free parameters, which were estimated for each participant: three parameters of the perceptual model (JGET HGF; *ω*_*x*_, 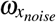, 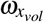), two parameters of the bucket position part of the action model (*σ*_*b*_, *σ*_*precision*_), and five parameters of the confidence part of the action model 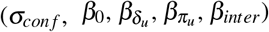. Supplementary Table S1 lists the fixed initial values used for the other belief states that are not estimated.

#### Priors for the cognitive model

##### Prior elicitation

For all parameters of the cognitive model, we specified Gaussian priors whenever possible. However, these distributions were truncated for two of the *ω* parameters (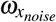 and 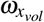) to avoid regions of the parameter space where the assumptions underlying the variational inversion of our perceptual model are violated. The prior means for the confidence regression model (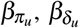 and *β*_*inter*_) were set to zero, to enable the model to capture any potential effect sizes that drive confidence formation. As *β*_0_ was intended to capture the confidence intercept for each participant, which tends to be non-zero and strictly positive, its mean was set to 0.6 (and not 0.5 because the empirical population confidence average is higher than 0.5, see Fig. 1b). In Fig. 3, the prior densities for each free model parameter are shown, while the exact definition for the prior distributions for each free parameter can be found in Table S2 of the Supplementary Materials.

**Figure 3.**
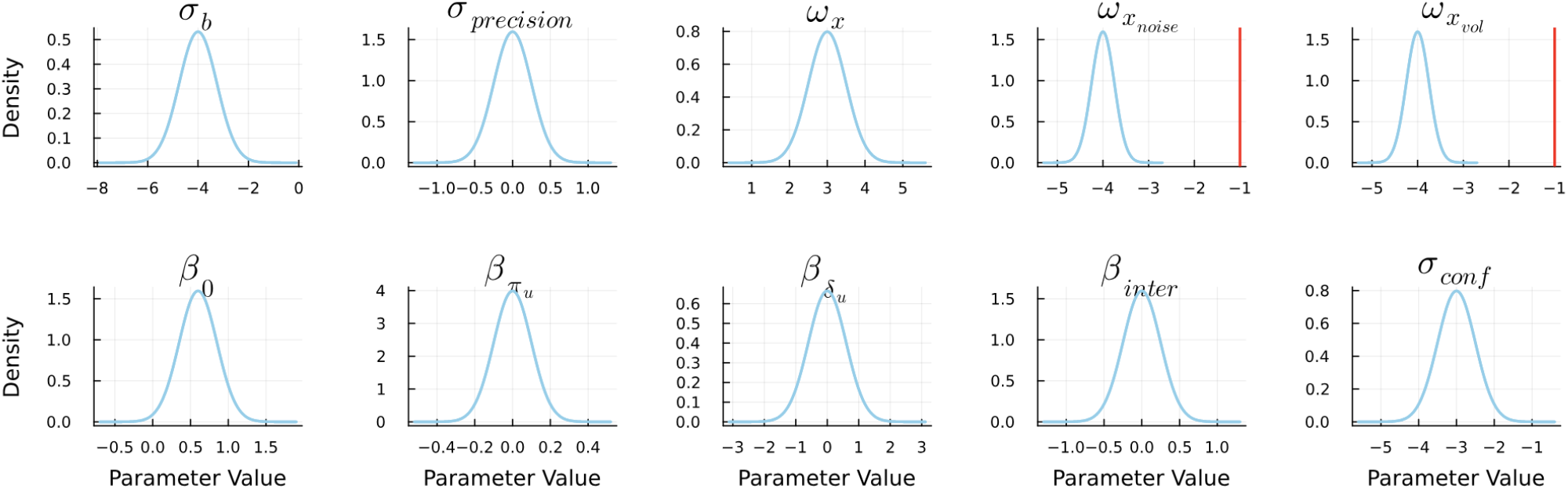
Prior densities for each free parameter for our cognitive computational model. Truncation of the distributions for 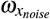 and 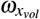 is marked with a red vertical line.

##### Prior predictive checks

Prior selection was tuned and qualitatively assessed by generating and visualising simulated model responses obtained from 75 random draws from the prior. Supplementary Fig. S1 displays simulated model responses (simulated predictions 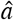 and confidence ratings 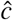) for the particle landing position trajectories (*U*_*t*_) of three participants (392, 307, 233) from our data set. For visualisation, we display the prior predictive density together with the responses of the participant. Note that these traces were generated by applying the model (with a parameter set drawn from the prior) to the traces that were also provided to the participants.

#### Model inversion and validation of computation

The HGF was implemented using the `HierarchicalGaussianFiltering.jl` library (v0.7.0), and extended to a full cognitive model using the `ActionModels.jl` (v0.7.3) sister-library. We fitted the cognitive model to the behavioural data using a variant of Markov chain Monte Carlo (MCMC)^41^ sampling. Specifically, we relied on the No U-Turn Sampler (NUTS)^42^ implemented as part of the `Turing.jl`^43,44^(v0.37.0) library. For each participant, two chains of 1000 samples each were run to produce posterior estimates of parameter values. Visual inspections of chain traces and quantitative diagnostics (potential scale reduction factor 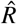 *<* 1.05) were used to check that the chains had converged and were well-mixed^45^. A second iteration with an extra chain and a more exhaustive sampler was used for those participants where the sampler had not converged during the first iteration. In total, the sampler converged 98.3% of the fitted participants (see “Convergence Checks” section in the Supplementary Materials and Fig. S4 for details). The remaining, non-converged participants were discarded from further analysis.

In addition, we performed a parameter recovery analysis^37,46^to confirm that parameter estimates can reflect actual data-generating mechanisms. We generated a synthetic dataset by drawing 30 parameter values from the prior for all free parameters of our cognitive model (‘prior noise’ condition). We repeated this procedure twice to assess parameter recovery for different noise conditions consisting of: a ‘posterior noise’ condition where both baseline action noise (*σ*_*b*_) and confidence noise (*σ*_*con f*_) parameter values were sampled from *N* (*−*1, 0.75), representing estimated empirical noise levels; a ‘high noise’ condition where *σ*_*b*_ and *σ*_*con f*_ values were both drawn from *N* (1, 0.75). In each noise condition, the sampled parameter values were used to simulate synthetic behaviour (bucket positions *a* and confidence ratings *c*), using the empirical input sequences (particle landing positions *U*_*t*_) from a randomly selected participant 53. We then fitted the model to the synthetic behavioural data using the same settings as for model inversion on the empirical dataset. Pearson’s correlation coefficients (*r*) between simulated and estimated parameter values were used to quantify parameter recoverability for all three noise conditions. Excellent recoverability was observed in the ‘prior noise’ condition (*r ≥* 0.9), good recoverability for the ‘posterior noise’ condition (*r ≥* 0.6), and bad to moderate recoverability of the ‘high noise’ condition (0 *≤ r ≤* 0.8). Visualisations as well as a more in depth discussion of the results from the parameter recovery analysis are presented in the Parameter Recovery Analysis section of the Supplementary Materials.

#### Model evaluation

Following model inversion on the empirical data set (N=430), we performed posterior predictive checks comparing the empirical responses and model fits for each participant using the posterior median values, as well as 30 random posterior samples. Specifically, the median from the single-participant posterior distributions, as well as the posterior samples, were used to generate simulated behavioural responses. These simulations were visually compared to empirically observed responses for six selected participants representative of different levels of goodness of fit for the confidence ratings. In addition, we examined the quality of our model fits using different metrics. First, we calculated Pearson’s correlation coefficients (*r*) between predicted (using posterior median values) and observed responses (bucket placements *a* and confidence ratings *c*). Second, model fit residuals for confidence ratings were calculated and visualised. Third, we compared empirical and model-based confidence ratings around experimental change-points (i.e. changes in the mean of the observation generating distribution). As a basis for interpretation of our results obtained from model inversion on the empirical dataset, we visualised single-participant as well as group-level posterior parameter estimates and contrasted these with our specified prior distributions. In addition, for each parameter, we extracted the posterior samples for every participant and computed the posterior mean and 80% credible interval (10%–90% quantiles). Participants were then ordered by their posterior median for that parameter. We visualized the resulting subject-level estimates using forest-style plots, where points represent posterior means and horizontal lines indicate the 80% credible intervals. To formally test whether parameter distributions systematically deviated from zero, we performed a sign-permutation test (10, 000 iterations). For each parameter, we calculated the empirical proportion of participants whose 80% credible interval fell entirely to one side of zero (e.g., lower bound *>* 0 or upper bound *<* 0). Under the null hypothesis of a symmetric distribution centered at zero, we repeatedly generated a null distribution by randomly flipping the signs of each participant’s posterior chain with equal probability (*p* = 0.5). The empirical *p* value was computed as the proportion of null iterations yielding a proportion of directed credible intervals equal to or greater than the observed data.

### Relation to transdiagnostic factor scores

To assess the relationship between the participant-specific cognitive model-derived parameter estimates and the transdiagnostic factor scores (AD, CIT, and SW), we performed linear regression analysis, accounting for potential confounding of demographic variables (age, gender, and IQ). We used linear regression models (from GLM.jl v1.9.0) of the form

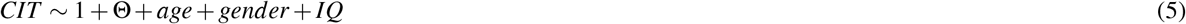

where Θ represents a cognitive model parameter. This analysis was restricted to the CIT score and the parameters related to the metacognitive part of our model 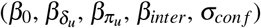. We therefore used Bonferroni-corrected significance level 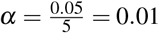.

### Code availability

The code needed to run the analysis, as well as a curated version of the original dataset, can be found in: https://gitlab.ethz.ch/tnu/code/felezetal_*c*_

## Results

### Model Evaluation

Results from posterior predictive checking are presented in Fig. 4 and Supplemantary Fig. S6. Specifically, Fig. 4 displays simulated and empirical confidence traces for six exemplary participants, ordered by an increasing confidence goodness-of-fit, as measured with the Pearson’s correlation coefficient (*r*) between the empirical and the simulated response generated with the median of the posterior. In general, the simulated responses change more gradually compared to the empirical traces. This is observed even for cases where the *r* is relatively high.

**Figure 4.**
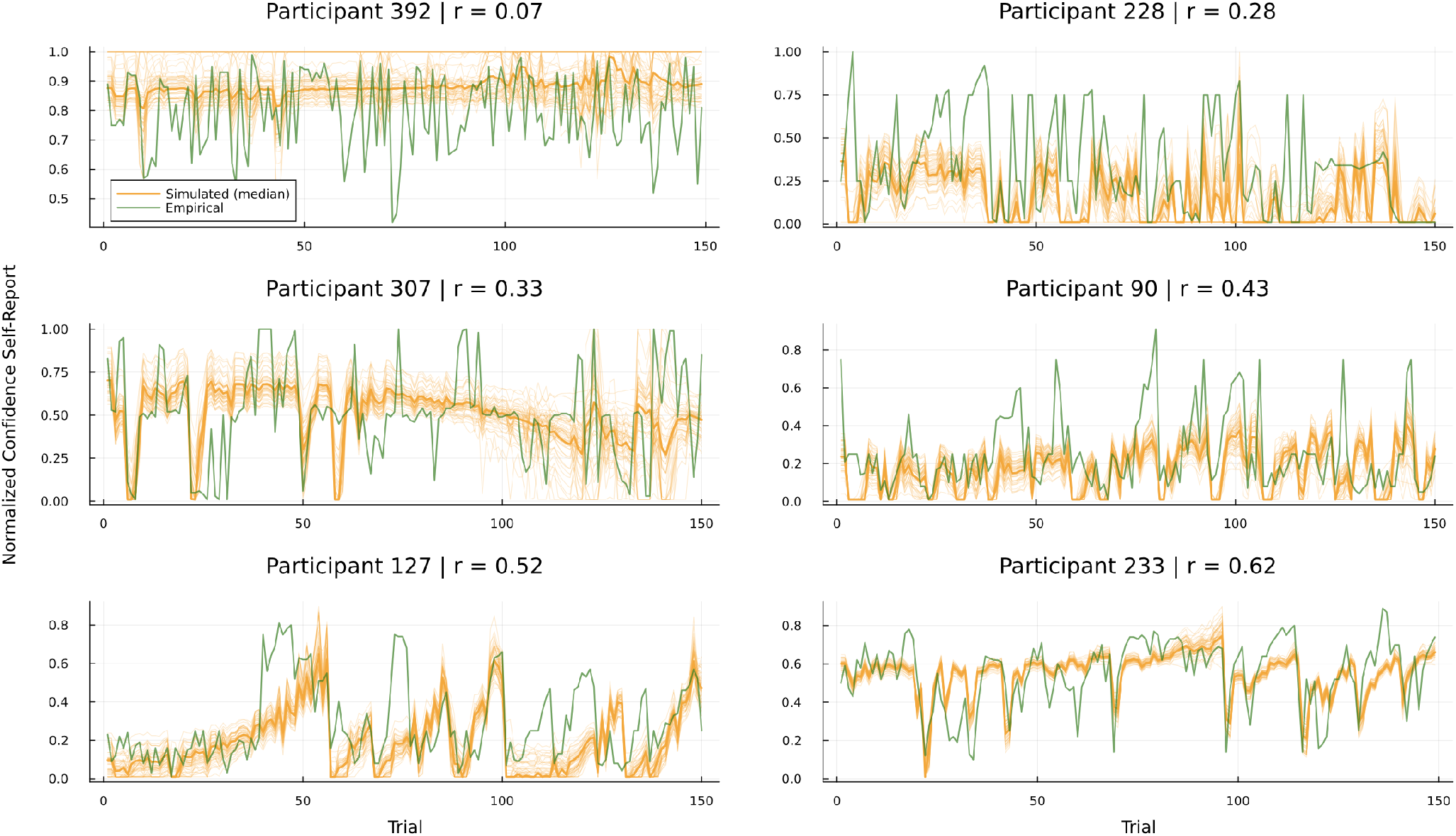
Simulated (orange) vs empirical (green) confidence responses from six participants, ranging from bad to good goodness-of-fit. The thick orange line identifies the simulated response from the median of the approximate posterior, while the thinner orange lines show the behaviour generated from the 30 random samples.

A histogram of *r* values representing the confidence rating goodness-of-fit is displayed in Fig. 5a. The white dot represents the average *r* across participants 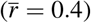. The confidence goodness-of-fit was negatively correlated with the CIT score (*r* = *−*0.33, 95%*CI*[*−*0.41, *−*0.25], *p <* 1*e −*11), but not with the other transdiagnostic dimensions (AD and SW). In Fig. 5b, we show the residual distribution for the confidence modelling, aggregated across all participants of the empirical dataset. The residual distribution is centered around zero, suggesting no systematic under- or overestimation of confidence ratings by our model. The single-participant residual plot is shown in Supplementary Fig. S9. Figure 5c displays simulated and empirical confidence ratings around experimental change-points. Model-based predicted confidence ratings accurately capture the sharp decline and consequent recovery in confidence ratings in the five trials following an experimental change-point.

**Figure 5.**
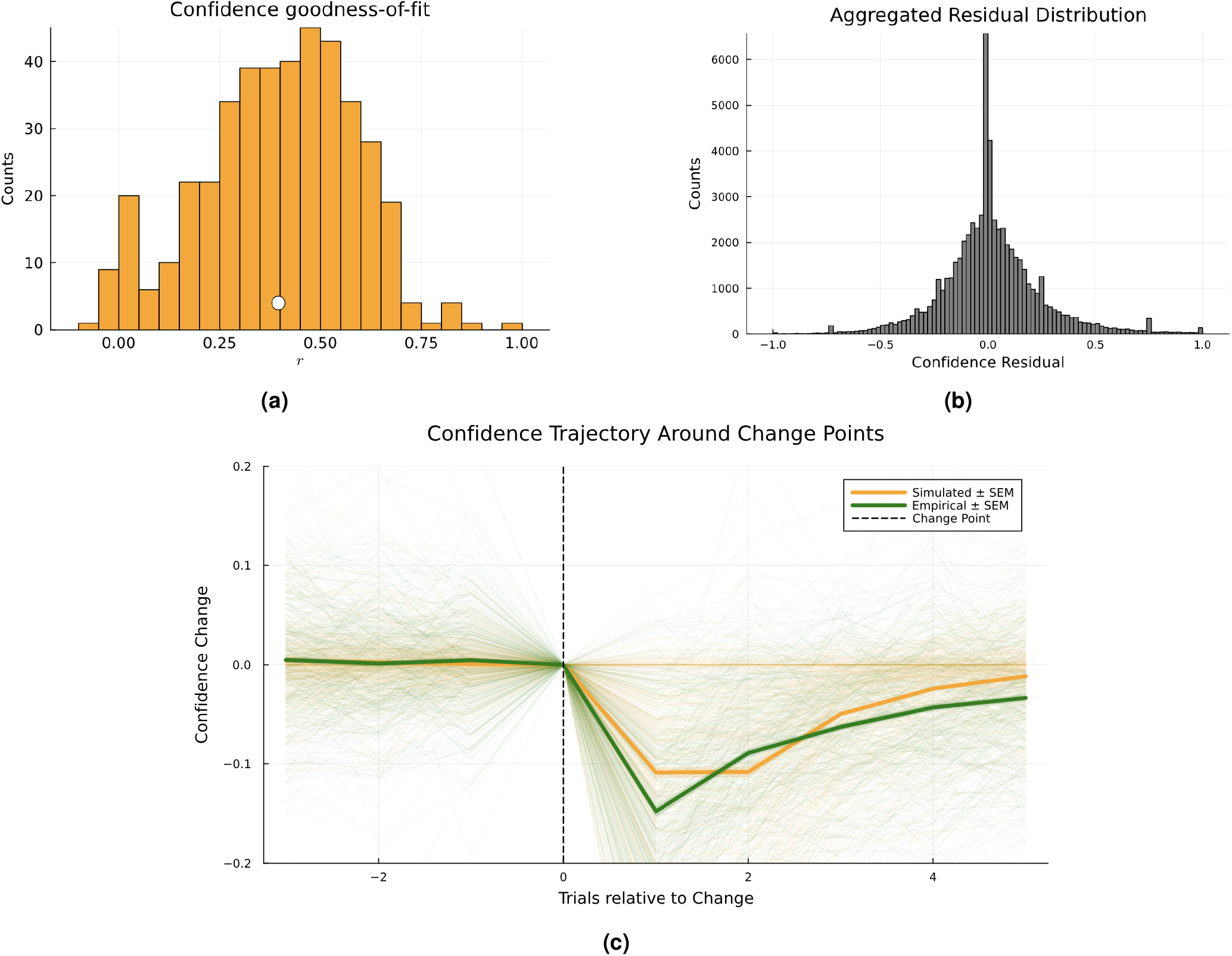
(a) Histogram of confidence rating goodness-of-fit, as measured by Pearson’s correlation coefficient (*r*). The white dot represents the average *r* across participants (*r* = 0.4). (b) Residual distribution for the confidence modelling. (c) Aggregated confidence trajectories (simulated vs empirical) around experimental change-points, across all participants and change-points. The thin line captures the single participant confidence dynamics.

Supplementary Fig. S5 displays a histogram showing goodness-of-fit as measured by the *r* between empirical and simulated bucket positions (*a*). For bucket positions, model fit is excellent on average 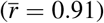 and does not correlate with any transdiagnostic factor score. In Supplemantary Fig. S6, we show the predicted model responses for the bucket placements (*a*), together with the empirical responses.

### Posterior Parameter Estimates

Figure 6 displays the subject-level posterior mean estimates, with the 80% credible intervals. The estimated group mean for 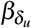 was negative (group mean = -0.2863), indicating a negative influence of prediction error magnitude (related to particle landing positions) on confidence ratings. On an individual level, the estimated effect was negative without overlapping with 0 (using an 80% credible interval) for 55.3% of participants (*p <* 0.001, sign-permutation test), suggesting that the negative group mean reflects a general negative effect among participants. In contrast, the estimated group mean 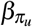 was positive (group mean = 0.0166), with 50.9% of the participants having individually positive effects with 80% credible intervals not overlapping with 0 (*p <* 0.001, sign-permutation test), showing a positive influence of precise predictions on confidence ratings. The posterior group mean for *β*_*inter*_ became negative (group mean = -0.0863, *p <* 0.001, sign-permutation test), with 37.2% of the participants showing this shift. Finally, the shift in negative direction by the group mean posterior of *σ*_*precision*_ (group mean = -0.2259) suggests a negative influence of the input prediction precision on bucket position action noise; meaning that the lower the precision (or the higher the variance) of the agent’s prediction about the particle landing position, the noisier becomes its bucket placement. This shift was present in 81.1% of the participants (*p <* 0.001, sign-permutation test). Supplementary Fig. S7 displays histograms of the subject-level posterior medians (i.e., parameter estimates used for the transdiagnostic score analysis) in green, as well as the group-level posterior means in blue, with the prior distributions overlaid in red for reference. It should be noted that the model estimates a higher posterior noise both for the bucket positions and for the confidence ratings compared to the chosen prior, as seen in the first two subplots for *σ*_*con f*_ and *σ*_*b*_.

**Figure 6.**
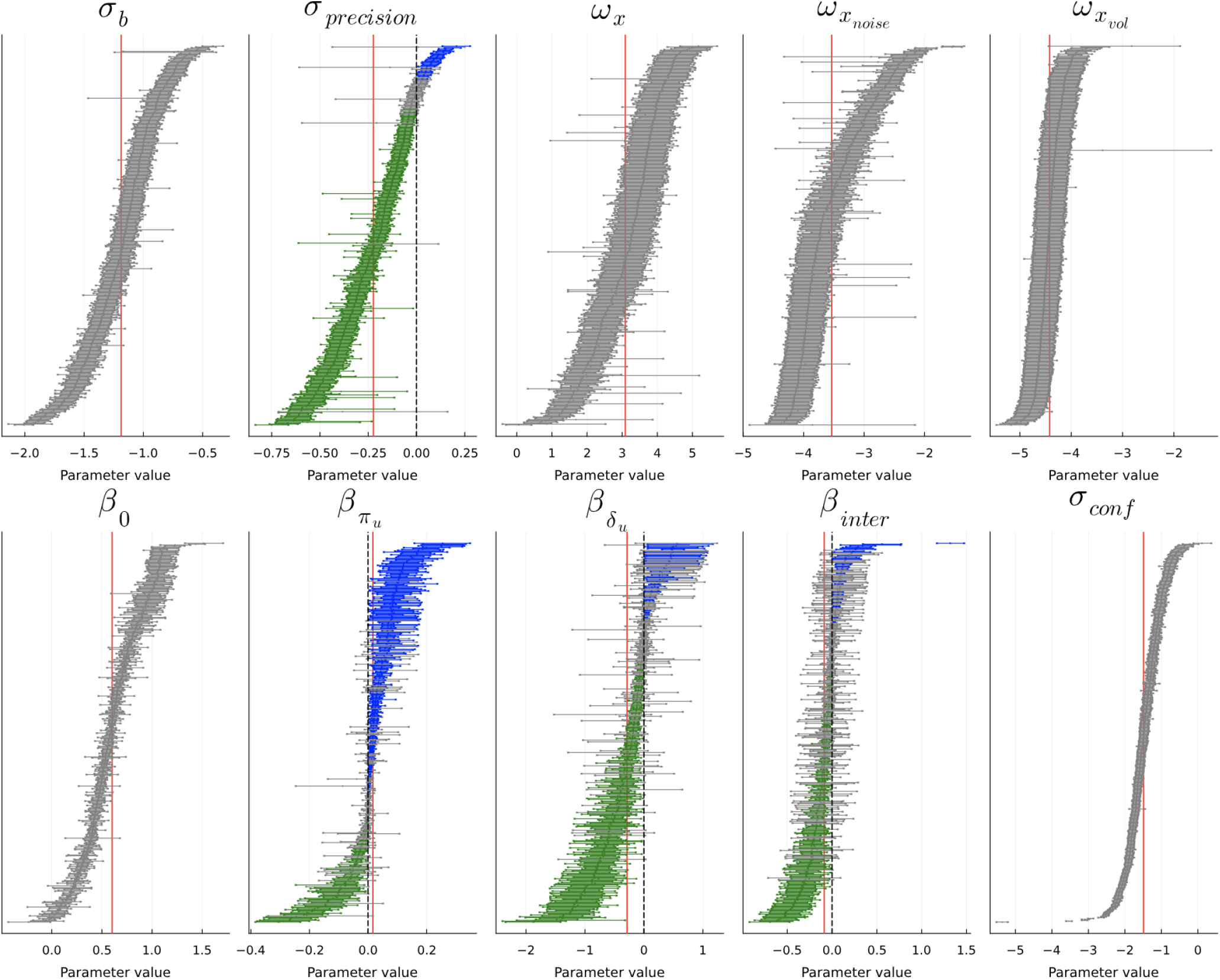
Subject-level posterior estimates for all model parameters. Each dot represents the posterior mean for a participant, and horizontal lines indicate the 80% credible interval (10th–90th percentile). Participants are ordered by increasing posterior median for each parameter. For the parameters with 0-mean priors, we show in green the participants for which the posterior mean (with the 80% credible interval) is strictly negative, blue if positive, and gray when the credible interval includes zero. Red vertical lines indicate the group mean for each parameter.

### Relation to Transdiagnostic Factor Scores

Linear regression analysis of CIT scores and cognitive model parameter Θ 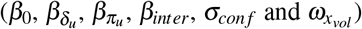, where age, gender and IQ were included as confounding variables, yielded the significant regression coefficients for parameters *β*_0_ and 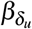 (Bonferroni-corrected *α* = 0.0083). Detailed results of the regression analysis are provided in Table 1 and scatter plots of CIT scores and parameters *β*_0_ and 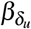 are shown in Supplementary Fig. S10.

**Table 1.** Results for the linear regression analysis.

| $\Theta$ | Regression Coef. | CI 95% | p value | BC significant ($\alpha = 0.01$) |
| --- | --- | --- | --- | --- |
| $\beta_0$ | 0.89 | 0.57, 1.21 | $< 1e-7$ | Yes |
| $\beta_{\delta_u}$ | 0.52 | 0.31, 0.72 | $< 1e-6$ | Yes |
| $\beta_{\pi_u}$ | 1.18 | 0.05, 2.31 | 0.0397 | No |
| $\beta_{inter}$ | 0.66 | 0.15, 1.17 | 0.0104 | No |
| $\sigma_{conf}$ | -0.21 | -0.38, -0.03 | 0.022 | No |

The significant positive regression coefficient for *β*_0_ indicates that, after controlling for age, gender, and IQ, higher CIT scores were associated with higher values of *β*_0_. In our cognitive computational model, *β*_0_ captures the baseline level of confidence ratings (Eq. 4), corroborating an overconfidence present in individuals with higher CIT scores^12^. We also observed a significant positive regression coefficient for 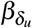 . This parameter weighs the influence of the magnitude of the prediction error about sensory input on confidence ratings. Given that most participant-level posterior medians for 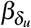 were negative, the positive regression coefficient suggests that individuals with higher CIT scores have a less negative coupling between prediction errors and confidence compared with individuals with lower CIT scores. These findings are consistent with previous findings in the literature^12^.

## Discussion

In this study, we introduced a cognitive computational model for learning and metacognition in the flying particle predictive inference task. We used a JGET HGF in combination with a multivariate action model to jointly model predictions (bucket positions *a*) and confidence ratings (*c*) based on a sequence of sensory inputs (particle landing positions *U*). We provided a detailed characterisation of our model following the general steps of Bayesian workflow for applied Bayesian modelling^37,39^ using both simulations and inference on a publicly available dataset (N=430) from the predictive inference task^12^. Model evaluation revealed that our model was able to achieve excellent fit for prediction responses (bucket positions *a*) and moderate to good fit for confidence rating data (*c*). Examination and quantification of posterior parameter estimates at the group level using 80% credible intervals revealed a negative effect of prediction error magnitude (related to sensory input; group mean = -0.2838) and a positive effect of prediction precision on confidence (group mean = 0.0163). Linear regression analysis between parameter estimates of our cognitive computational model and transdiagnostic factor scores derived from psychological questionnaire data revealed a significant positive associations between *β*_0_ as well as 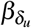 and CIT scores (Bonferroni-corrected), which aligns with previous literature findings^11,12^.

### A Cognitive Computational Model of Learning and Metacognition

Our joint prediction-confidence model was based on the HGF framework. We designed an action model capable of generating not only bucket position predictions but also confidence reports based on inferred belief states using the HGF. For the latter part, we built a general linear model of confidence ratings with four regressors: *β*_0_ captured baseline confidence levels, 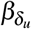 weighted the influence of the PE about sensory input from the previous trial on reported confidence (PE-confidence coupling), 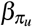 weighted the contribution of the precision of the sensory input prediction to confidence ratings, and *β*_*inter*_ scaled the effect of the interaction between PE and precision on confidence ratings. Hence, our model is incorporates the widely discussed feedback effect in confidence reports in the literature^47–50^, in which external performance feedback systematically alters subsequent confidence judgments. In relation to our model for predictive inference task data, the PE represents a combination of external performance feedback (observed outcomes) and internal expectations (predicted outcome), which is consistent with Bayesian accounts of brain function such as predictive coding. By using the prediction precision, our model aligns with the view of confidence as reflecting the Bayesian posterior probability of a choice being correct^13–16,51^.

Our proposed model was inverted using a variant of MCMC for every participant. We found a good average confidence goodness-of-fit, as measured by the average 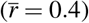, highlighting the ability of our model to capture confidence dynamics, while maintaining an overall excellent fit for the prediction responses 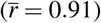. The residual analysis further indicates that there is no systematic bias in the predicted confidence ratings by our model (Fig. 5b). The change-point analysis (Fig. 5c) demonstrated that the model captured both the immediate decrease in empirical confidence ratings after the change-point as well as the gradual increase back to baseline in the following trials. Qualitative and quantitative analysis of the approximate posterior distributions revealed a shift in negative direction of the 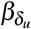 posterior, which was centered around zero. In other words, confidence ratings were lower in trials where prediction error magnitude was higher. Similarly, the group mean for 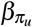 was positive (0.0163), suggesting a positive influence of sensory input prediction precision on confidence ratings.

Interestingly, confidence goodness-of-fit (Fig. 5a) correlated negatively with CIT scores, suggesting that our model captures confidence dynamics more accurately in individuals with lower CIT scores. This corroborates a finding from previous work examining the same dataset that decoupling between confidence ratings and task performance increases with higher compulsivity scores^12^. A possible explanation for this finding could be that self-evaluation in individuals with high CIT scores is more noisy. Alternatively, it has been suggested that confidence in highly compulsive individuals arises through different forms of information processing that are currently not captured by our model, such as increased emotional processing^52^.

### Prediction, confidence, and transdiagnostic factor scores

When regressing the model-derived parameters from the predictive inference task to the CIT transdiagnostic factor score, we found a significant positive regression coefficient for *β*_0_ in relation to CIT scores (*β* = 0.89, 95% CI [0.57, 1.21], *p <* 1*e* − 7). This is in line with previous work using the same task that found that compulsivity was linked to inflated confidence^11,12^.

However, the finding of inflated confidence in compulsivity in learning tasks contrasts with previous findings of decreased confidence in patients with OCD performing memory tasks^3^. The discrepancy between increased confidence in learning and decreased confidence in memory in compulsive individuals could be due to several reasons. First, there is evidence for domain-specific differences in metacognition^53^, such that an individual’s metacognitive sensitivity or confidence calibration in one cognitive domain (e.g. learning) does not necessarily generalise to other domains (e.g. memory, perception). Second, previous studies did not account for potential comorbidities present in patients (see Ref.^3^ and references therein). Since depression has been associated with a general decrease in confidence^54^, the co-occurrence of depressive and compulsive symptoms could mask the underlying direction of the association between compulsivity and confidence. Our analysis overcomes this problem by adopting the approach introduced in Ref.^38^ which makes use of transdiagnostic factor scores^12,55^ as opposed to raw questionnaire data.

Moreover, we found a significant positive regression coefficient for 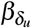 in relation to CIT scores (*β* = 0.52, 95% CI [0.31, 0.72], *p <* 1*e −* 6). Our model may provide a more nuanced perspective on action-confidence coupling in the predictive inference task, i.e., the size of action updates have been shown to be related to confidence ratings within-subjects, “such that lower confidence was linked to larger updates”^12^. Vaghi et al.^11^ initially found a reduced coupling between actions and confidence ratings in OCD patients, which was positively correlated with OCD symptoms severity. In Ref.^12^, this finding was replicated in a general population sample and the authors further demonstrated that a reduction in action–confidence coupling was not specific to any individual questionnaire score but could instead be accounted for by a single transdiagnostic dimension (the CIT score). Our approach frames these results as a decoupling between PE and confidence, providing a Bayesian brain perspective on compulsivity.

### Limitations and Future Work

There are several limitations that deserve to be mentioned. First, our suggested trial-by-trial computational cognitive model of confidence rating data consisted of a simple linear regression with four coefficients, which represents a descriptive but not mechanistic account of confidence formation, even though some mechanistic components are present in the generative model of the HGF. This reflects a deeper issue in the scientific literature on confidence formation, namely, the current lack of a well-established theory of confidence formation in the context of learning. This makes the development of mechanistic models of confidence formation particularly challenging but at the same time highlights the need for these types of models to advance our understanding of the matter. By definition, confidence ratings reflect subjective, introspective experiences and may therefore integrate multiple underlying factors. Future work may build on our model and combine it with ideas of action-dependent models, in which confidence ratings directly reflect the probability of being correct^2,3,16^, instead of our indirect approach using prediction precision.

Second, while our model is able to capture meaningful variations in confidence ratings, there is still room for improvement, as indicated by the many moderate fits. Our current approach provides a very simple formalisation of confidence ratings as a linear combination of PE, precision of the sensory input prediction, as well as their linear interaction. It is likely that confidence formation is a more complicated process, including non-linear relations as well as dependence on other quantities such as simulated actions akin to policy roll-outs in models of action selection such as reinforcement learning^56^ or active inference^57^. As seen in the posterior predictive checks (Fig. 4), the fast changes in confidence are not well captured by our model. Bayesian model comparison represents one potential tool to differentiate between alternative hypotheses in the form of different computational models^37,58^.

An important methodological choice in our action model concerned the response function of the latent confidence expectation *η*_*c,t*_, which restricted confidence values to the open interval (0, 1) through a piecewise linear function, preventing it from reaching the boundary values that would otherwise render the truncated normal likelihood numerically degenerate. While this approach proved effective and computationally simple, several alternative strategies exist for handling the bounded nature of confidence ratings in a less aggressive way. A logit or probit link function, for instance, would map an unconstrained linear predictor onto (0, 1) without requiring explicit clamping, though at the cost of less directly interpretable regression coefficients.

Alternatively, a Beta distribution parameterised in terms of mean and precision could be used as an observation model for confidence ratings, which naturally respects their bounded support. We opted for parameter clamping over these alternatives primarily to preserve the direct interpretability of the linear combination of regressors in *η*_*c,t*_ (Eq. 4) while ensuring stable model inversion; future work could nonetheless investigate whether these alternative link functions or observation models yield improved fit or a more principled treatment of the bounded nature of confidence data.

A different direction for future research could be to specify a hierarchical model of task data and questionnaire measures. This would enable joint inference on latent variables underlying behaviour as well as their relation to questionnaire data, which is informationally superior to estimating parameters and then relating them to the questionnaire data in a separate analysis. However, it should be noted that, although there are statistical benefits to accounting for multiple sources of uncertainty together, the resulting inversion problem becomes more challenging to handle in practice.

The last limitation pertains to the dataset we used for our analysis. Each participant was exposed to a unique input sequence, meaning that comparing the inferred parameters relies on an assumption, as these parameters are conditionally dependent on the specific inputs. Despite its considerable size (N=437), it still represents one specific instantiation of the predictive inference task. Hence, future work should assess the generalisability of our results by applying our model to data from similar tasks such as the ones presented in Ref.^26^. In addition, the dataset represents a cross-sectional sample of the general population. While this is extremely useful and necessary for the development of cognitive computational models of task behaviour, it merely provides a first basis for hypothesis generation related to potential disease mechanisms underlying mental health disorders. Testing these hypotheses, such as a potential metacognitive deficit underlying compulsive behaviour and potentially intervening on causal links requires longitudinal studies with patient populations.

## Conclusion

In this work, we presented an HGF-based trial-by-trial model of prediction responses and confidence ratings in a predictive inference task. By applying it to an empirical dataset from Ref.^12^, we demonstrated that our model captured significant variation in prediction responses and confidence ratings across a variety of different metrics. In addition, we found evidence for a negative effect of sensory input prediction error magnitude on confidence ratings and a positive effect of prediction precision on confidence ratings, respectively. Furthermore, we replicated previous findings using data from the same task that describe associations between compulsivity and inflated confidence^11,12^. In this work, the previously observed decoupling between action updates and confidence in individuals with high CIT scores is reframed as a dissociation between prediction errors and confidence ratings. Together, the results demonstrate the robustness of our methodology for combined prediction-confidence fitting, which has great potential to characterise individual differences in metacognitive belief formation that can lead to novel psychopathological insights.

## Acknowledgements

The project that gave rise to these results received the support of a fellowship from the “la Caixa” Foundation (ID 100010434). The fellowship code is B006495. We thank Klaas Enno Stephan and Jakob Siemerkus for their guidance and helpful discussions during the development of this work.

## Author contributions statement

E.F. conceived part of the model development, implemented the software, conducted the analysis, and wrote the manuscript.

A.J.H. conceived and supervised the model development and analysis. P.T.W. and J.H. provided feedback throughout the work.

P.T.W. performed a code review. All authors reviewed and edited the manuscript.

## Additional information

### Accession codes

TBD

