## Supplementary Materials for "Modelling Metacognition: A Joint Prediction-Confidence Model for Predictive Inference Task Data"

**Table S1.** Values of fixed parameters for the HGF (perceptual model). The parameters not shown here were set to their default value.

| | | |
|---|---|---|
| $\mu_{x_0}$ | initial mean of $X$ | 0 |
| $\pi_{x_0}$ | initial precision of $X$ | 0.25 |
| $\mu_{x_{noise_0}}$ | initial mean of $X_{noise}$ | 10 |
| $\pi_{x_{noise_0}}$ | initial precision of $X_{noise}$ | 0.33 |

List of free parameters of the cognitive model:

- $\omega_x$: tonic volatility for state $X$.
- $\omega_{x_{noise}}$: tonic volatility for state $X_{noise}$.
- $\omega_{x_{vol}}$: tonic environmental volatility for the upper layer state $X_{vol}$.
- $\sigma_b$: baseline action noise.
- $\sigma_{precision}$: weight of influence of the input prediction precision on the noise.
- $\sigma_{conf}$: baseline confidence noise.
- $\beta_0$: baseline confidence level.
- $\beta_{\delta_u}$: influence of the absolute prediction error from the previous trial on the confidence rating.
- $\beta_{\pi_u}$: influence of the prediction precision on the confidence rating.
- $\beta_{inter}$: influence of the interaction between input prediction precision and absolute prediction error from the previous trial on the confidence rating.

**Table S2.** Prior distributions for perceptual and action model parameters used for model inversion. $\mathcal{N}(\mu,\sigma^2)$ and $\mathcal{TN}(\mu,\sigma^2\ ;\text{min, max})$ stand for Normal (with $\mu$ mean and $\sigma^2$ variance) and Truncated Normal (with $\mu$ mean and $\sigma^2$ variance, and bounded in min, max range), respectively.

| | |
|---|---|
| $\omega_x$ | $\mathcal{N}(3,0.5)$ |
| $\omega_{x_{vol}}$ | $\mathcal{TN}(-4,0.25;-\infty,-1)$ |
| $\omega_{x_{noise}}$ | $\mathcal{TN}(-4,0.25;-\infty,-1)$ |
| $\sigma_b$ | $\mathcal{N}(-4,0.75)$ |
| $\sigma_{precision}$ | $\mathcal{N}(0,0.25)$ |
| $\sigma_{conf}$ | $\mathcal{N}(-3,0.5)$ |
| $\beta_0$ | $\mathcal{N}(0.6,0.25)$ |
| $\beta_\pi$ | $\mathcal{N}(0,0.1)$ |
| $\beta_{\delta_u}$ | $\mathcal{N}(0,0.6)$ |
| $\beta_{inter}$ | $\mathcal{N}(0,0.25)$ |

#### HGF generative model equations

Formally, in the HGF's generative model, each continuous state node $n$ in the HGF network follows a Gaussian random walk where the volatility depends on any volatility parents $j \in I$. In our case, the value of the state node $x_t$ at trial $t$ is given by:

$$X_t \sim \mathcal{N}(X_{t-1}, \exp(\omega_x + X_{vol,t-1})) \tag{6}$$

The input node is then sampled from a Gaussian distribution with mean and variance defined by its value parent $i$ and noise parent $j$, respectively:

$$u_t \sim \mathcal{N}(X_t, \exp(X_{j,t})) \tag{7}$$

The higher-level state $X_{vol}$ is modelled as a Gaussian random walk with tonic environmental volatility $\omega_{x_{vol}}$:

$$X_{vol,t} \sim \mathcal{N}(X_{vol,t-1}, \exp(\omega_{x_{vol}})) \tag{8}$$

Inverting this generative model with a variational message passing scheme yields a belief updating algorithm with three steps: a prediction step, a belief update step, and a prediction error (PE) step. This algorithm is what is used as perceptual model when the HGF is used as cognitive model. The update equations are presented in the next section.

### HGF update equations

#### *Continous State Node update equations*

Here, we briefly summarise the general update equations used in the Hierarchical Gaussian Filter. See Ref.[25] for a full treatment. Note that several parameters, such as the autoconnection strengths and drift, are in the analysis for this article set to defaults so that they have no effect on the belief updating.

The prediction mean for timestep $t$ is the sum of the previous belief $\mu_{n,t-1}$, weighted by the autoconnection strength $\lambda_n$, and the predicted total drift $\mathrm{P}_{n,t}$:

$$\hat{\mu}_{n,t} = \lambda_n \mu_{n,t-1} + \tau_t \mathrm{P}_{n,t} \tag{9}$$

The predicted total drift is the sum of the tonic drift $\rho_n$ and the belief $\mu_{i,t}$ of each value parent $i$, transformed according to the non-linear transformation $g_{i,n}$ and weighted by the coupling strength $\kappa_{i,n}$ from that parent to the node $n$, all multiplied by the timestep size $\tau_t$:

$$\mathrm{P}_{n,t} = \rho_n + \sum_{i=1}^{N_{vapa}} \kappa_{i,n} g_{i,n}(\mu_{i,t-t}; \theta_{i,n}) \tag{10}$$

The precision of the prediction $\hat{\pi}_{n,t}$ is calculated as the inversion of the sum of the inverse precision of the previous belief $\pi_{n,t-1}$ and the total predicted volatility $\Omega_{n,t}$:

$$\hat{\pi}_{n,t} = \frac{1}{\frac{1}{\pi_{n,t-1}} + \tau_t \Omega_{n,t}} \tag{11}$$

The predicted total volatility is the sum of the tonic volatility $\omega_n$ and the belief $\mu_{j,n}$ of each volatility parent $j$, weighted by the coupling strength to that parent $\kappa_{j,n}$, and multiplied by the timestep size $\tau_t$:

$$\Omega_{n,t} = \exp\left(\omega_n + \sum_{j=1}^{N_{vopa}} \kappa_{j,n} \mu_{j,t-1}\right) \tag{12}$$

In the belief update step, the precision of the belief is calculated as the sum of the prediction precision $\hat{\pi}_{n,t}$, for each value child $i$ the precision of the prediction $\hat{\pi}_{n,t}$ weighted by the squared coupling strength $\kappa_{n,i}$, for each volatility child $j$ the sum of the precision prediction errors $\Delta_{j,t}$, weighted by the volatility-weighted precision $\gamma_{j,t}$ and the coupling strength $\kappa_{n,j}$, and for each probability child $k$ (i.e., binary state node value children), the inverse precision prediction $\frac{1}{\hat{\pi}_{k,t}}$ weighted by the squared coupling strength $\kappa^2_{n,k}$:

$$\begin{aligned}
\pi_{n,t} = \hat{\pi}_{n,t} + \\
&\sum_{i=1}^{N_{vach}} \left(\hat{\pi}_{i,t}\left(\kappa^2_{n,i} g'_{n,i}(\mu_{n,t-t}; \theta_{n,i}) - \kappa_{n,i} g''_{n,i}(\mu_{n,t-t}; \theta_{n,i})\delta_{i,t}\right)\right) + \\
&\sum_{j=1}^{N_{voch}} \left(\frac{1}{2}(\kappa_{n,j}\gamma_{j,t})^2 + \Delta_{j,t}(\kappa_{n,j}\gamma_{j,t})^2 - \frac{1}{2}\Delta_{j,t}\kappa^2_{n,j}\gamma_{j,t}\right) + \\
&\sum_{k=1}^{N_{probch}} \left(\kappa^2_{n,k}\frac{1}{\hat{\pi}_{k,t}}\right)
\end{aligned} \tag{13}$$

where $\gamma_{j,t}$ is the volatility-weighted prediction precision of the volatility child $j$:

$$\gamma_{j,t} := \Omega_{j,t}\hat{\pi}_{j,t} \tag{14}$$

and $\delta_i,t$ is the value prediction error of the child (equation 18). Note that when $g_{n,i}(x;\theta_{n,i})$ is the identity function $g(x) = x$, the term for children value simplifies to $\sum_{i=1}^{N_{vach}}\left(\hat{\pi}_{i,t}\kappa_{n,i}^2\right)$. The mean of the belief $\mu_{n,t}$ is then calculated as the sum of the mean of the prediction $\hat{\mu}_{n,t}$, for each value child $i$ (also including probability children) the precision-weighted prediction error $\varepsilon_{n,i,t}$ weighted by the coupling strength $\kappa_{n,i}$, and for each volatility child $j$ the precision-weighted precision prediction error $\mathscr{E}_{n,j,t}$ weighted by the coupling strength $\kappa_{j,n}$:

$$\mu_{n,t} = \hat{\mu}_{n,t} + \sum_{i=1}^{N_{vach}} \kappa_{n,i}\varepsilon_{n,i,t} + \sum_{j=1}^{N_{voch}} \kappa_{n,j}\mathscr{E}_{n,j,t} \tag{15}$$

The precision-weighted prediction error $\varepsilon_{n,i,t}$ between node $n$ and its value child $i$ is the child's value prediction error $\delta_{i,t}$ weighted by the precision weight $\psi_{n,t,t}$ and multiplied by the derivative of the non-linear transformation $g_{n,i}$ (which evaluates to 1 when using the identity transform):

$$\varepsilon_{n,i,t} = \psi_{n,i,t}g'_{n,i}(\mu_{n,t-t};\theta_{n,i})\delta_{i,t} \tag{16}$$

The precision weight $\psi_{n,i,t}$ is the relative precisions of $n$ and $i$:

$$\psi_{n,i,t} = \frac{\hat{\pi}_{i,t}}{\pi_{n,t}} \tag{17}$$

and the value prediction error $\delta_{i,t}$ for the child $i$ is the difference between its prediction and the posterior means:

$$\delta_{i,t} = \mu_{i,t} - \hat{\mu}_{i,t} \tag{18}$$

The precision-weighted precision prediction error $\mathscr{E}_{n,j,t}$ between node $n$ and its volatility child $j$ is then the child's precision prediction error $\Delta_{j,t}$ weighted by the precision weight $\Psi_{n,j,t}$:

$$\mathscr{E}_{n,j,t} = \Psi_{n,j,t}\Delta_{j,t} \tag{19}$$

The precision weight is the ratio of the child's volatility-weighted precision $\gamma_{j,t}$ and twice the parent's posterior precision $\pi_{n,t}$

$$\Psi_{n,j,t} = \frac{\gamma_{j,t}}{2\pi_{n,t}} \tag{20}$$

and the precision prediction error $\Delta_{j,t}$ of child $j$ is given as:

$$\Delta_{j,t} = \frac{\hat{\pi}_{j,t}}{\pi_{j,t}} + \hat{\pi}_{j,t}\delta_{j,t}^2 - 1 \tag{21}$$

Notably, these equations can be read as implement delta rule updates of the type used in classic learning rules from reinforcement learning (e.g., the Rescorla-Wagner), of the form $V_{t+1} = V_t + \alpha\delta$, where $V_t$ is the prediction, there is a prediction error $\delta$ for each child node, and the learning rate $\alpha$ depends on the precision weights and coupling strength variables. It is therefore possible to define the *implied learning rate* $\alpha_{n,i,t}$, that is, the learning rate that corresponds to using the HGF update equations on timestep $t$ for node $n$ and child $i$. For a value child $i$ or a volatility child $j$ the implied learning rate can be calculated as:

$$\begin{aligned} \alpha_{n,i,t} &= \kappa_{n,i}\psi_{n,i,t}g'_{n,i}(\mu_{n,t-t};\theta_{n,i}) \\ \alpha_{n,j,t} &= \kappa_{n,j}\Psi_{n,j,t} \end{aligned} \tag{22}$$

#### *Continuous Input Node update equations*

The continuous input varies from the continuous state node in some respects. The prediction mean for a continuous input node $U$ is the sum of the predictions $\hat{\mu}_{i,t}$ of its value parents $i$, weighted by coupling strengths $\kappa_{i,u}$. The bias $\rho_u$ is added as a constant value.

$$\hat{\mu}_{u,t} = \rho_u + \sum_{i=1}^{N_{vapa}} \kappa_{i,u}\hat{\mu}_{i,t-1} \tag{23}$$

The prediction precision is the inverse exponentiated sum of the previous posteriors of each noise parent $j$ and the tonic noise $\omega_u$:

$$\hat{\pi}_{u,t} = \frac{1}{\exp\left(\omega_u + \sum_{j=1}^{N_{vopa}} \kappa_{j,u}\mu_{j,t-1}\right)} \tag{24}$$

The posterior mean is equal to the input value $o_{u,t}$ for the input node $U$:

$$\mu_{u,t} = o_{u,t} \tag{25}$$

And the precision is infinite, since the value is known:

$$\pi_{u,t} = \infty \tag{26}$$

The value prediction error is calculated as usual, but the precision prediction error $\Delta_{u,t}$ used in the noise parents' updates uses the average posterior precisions $\bar{\pi}_{I,t}$ and average posterior means $\bar{\mu}_{I,t}$ from the value parents $i \in I$, instead of the node's own posterior mean and precision:

$$\Delta_{u,t} = \frac{\hat{\pi}_{u,t}}{\bar{\pi}_{I,t}} + \hat{\pi}_{u,t}(\mu_{u,t} - \bar{\mu}_{I,t})^2 - 1 \tag{27}$$

### Parameter Recovery Analysis

Pearson correlation coefficients ($r$) from parameter recovery analysis of our model are visualised in Fig. S2 for three different noise seetings: prior noise, posterior noise and high noise. In the prior noise setting, all of the ten free parameters of our model (three perceptual and seven response model parameters) show excellent recoverability when sampling from the priors with $r$ values between true and recovered parameters higher than 0.8 for all of the parameters. Under the posterior noise setting, recoverability decreases slightly, especially for parameters such as $\beta_{\delta_u}$, although still good recoverability levels are obtained ($r > 0.6$ in all cases, $r > 0.8$ for all cases except $\beta_{\delta_u}$). When increasing the noise even more (high noise setting), parameter recoverability decreases substantially for all parameters. Note however, that we did not find any of those noise levels in the posterior. In Supplementary Fig. S3, a scatter plot of the simulated (true) versus estimated (recovered) parameter values is displayed for all parameters and noise settings.

### Convergence Checks

In Fig. S4, the results averaged by the parameters and the results averaged by the participants are shown. The red line indicates the potential scale reduction factour boundary ($\hat{R} = 1.05$), above which which convergence is typically considered insufficient[59]. It can be seen that the mean $\hat{R}$ (± one standard deviation) falls below this threshold for every parameter (Fig. S4, left), while the participant-averaged $\hat{R}$ values for the vast majority of participants also remain below this limit (Fig. S4, right). Model inversion on the empirical dataset resulted in seven out of the 430 participants with an $\hat{R}$ value greater than 1.05. In other words, chain convergence was generally good across nearly all participants and parameters in the empirical dataset, although a few cases of poor convergence were observed. We were not able to identify a consistent pattern for these instances of poor convergence. As a consequence, we removed the participants for whom the sampler did not reach convergence during model inversion for further analysis.

The non-converged initial cases are participants: 192, 412, 291, 290, 88, 188, 31, 275, 113, 184, 138, 75, 283, 370, 25, 424, 379, 244, 437, 170, 380, 215, 238, 97, 143, 26, 61, and 66. The seven participants for which convergence cannot be assured after the second iteration were: 412, 188, 113, 75, 370, 25, and 437.

### Circular Data modelling

In the main text, we outlined how circular data from the predictive inference task can be preprocessed ('unrolled') to bring them into a form that is suitable for trial-by-trial computational cognitive modelling using HGF. Here, we describe two alternative approaches that could be used for circular data modelling using HGF. One approach would be to modify the HGF update equations (perceptual model) to explicitly handle circular data. Specifically, one would need to constrain the latent states to the 0–359º range and bound PEs between -180º and 180º. Although this approach could, in principle, provide a solution posed by the problem of circular data, it requires altering the underlying generative model of the HGF and deriving a new set of trial-by-trial update equations specifically designed for circular data.

Yet another approach would be to consider a different coding of the inputs (particle landing positions) and responses (bucket positions). Instead of using a one-dimensional input (angular position), every angular input and response would be transformed into a two-dimensional embedding using cosine and sine transformations. An HGF could be constructed with two input nodes and two state nodes naturally bounded from -1 to 1, which track the vertical and horizontal projection of the shooting direction, respectively. The $X_{noise}$ and $X_{vol}$ states could serve as shared parents for the two continuous state nodes of this HGF. While this approach provides another possible solution to the problem posed by circular data, it rests on a more complex perceptual model compared to the other two solutions. Moreover, the HGF was designed to provide a model belief updating in a participant's mind underlying behaviour, and we find it unrealistic to assume that participants track the two-dimensional angular embedding instead of just the angular shooting direction in the predictive inference task.

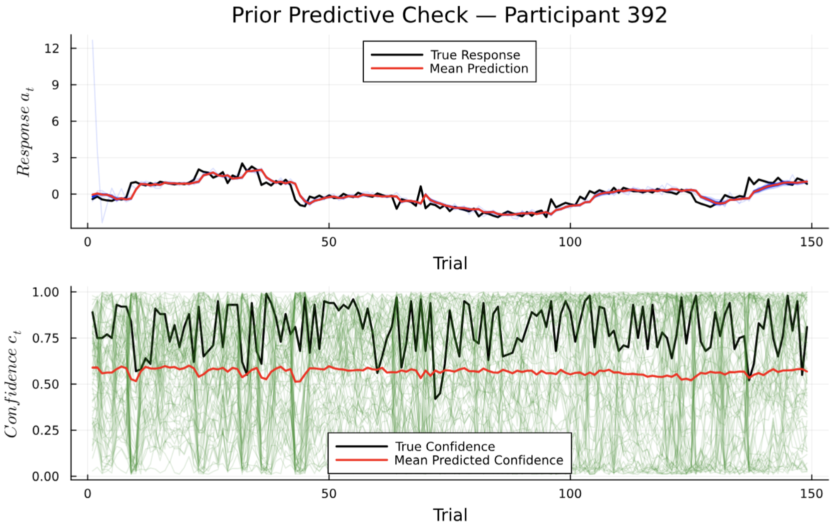


**(a)**

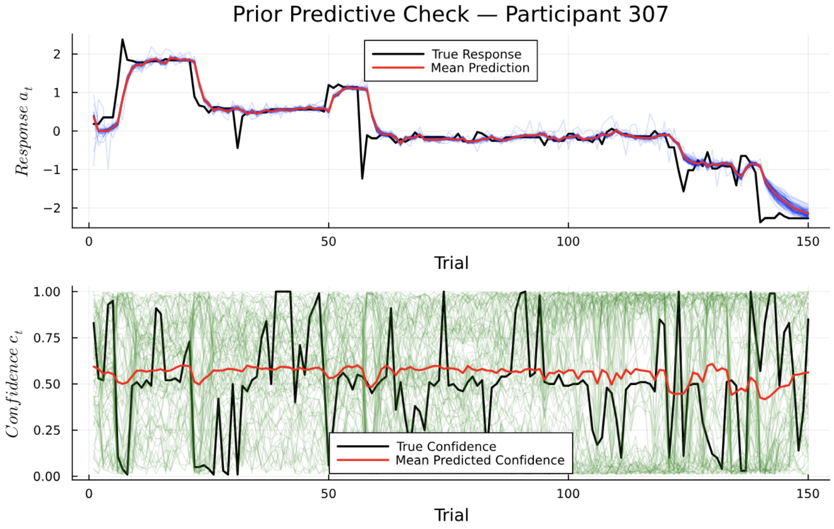


**(b)**

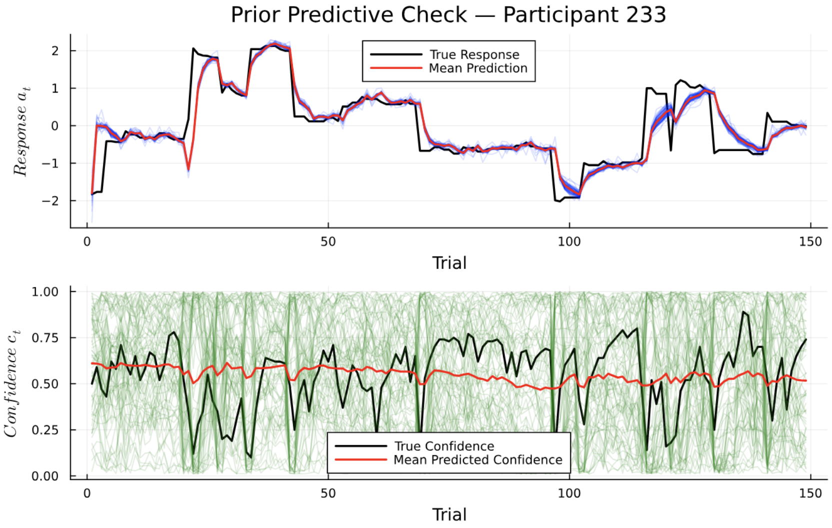


**(c)**

**Figure S1.** Prior predictive checks for participant 392 (a), participant 307 (b) and participant 233 (c). Red lines are used to identify the mean prediction and confidence response, while blue and green are used for individual responses obtained for one prior sampling, for the action predictions and confidence, respectively.

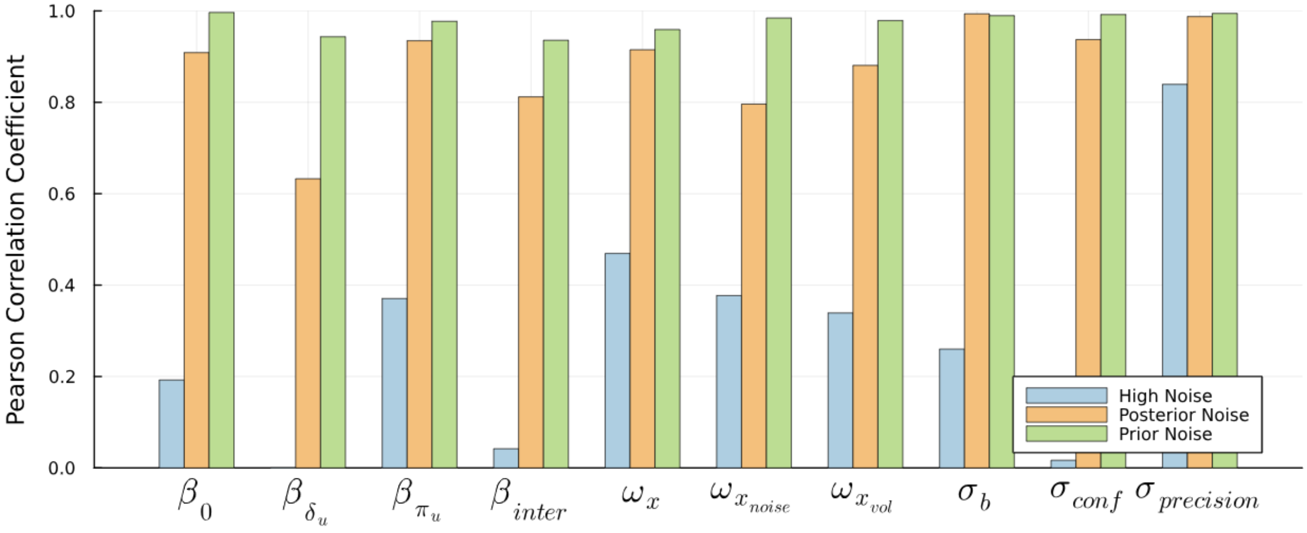


**Figure S2.** Pearson's correlation coefficients ($r$) from parameter recovery analysis for the prior, posterior and high noise case (green, orange and blue, respectively). Recoverability is good to excellent when sampling from the priors, but decreases with the increasing noise.

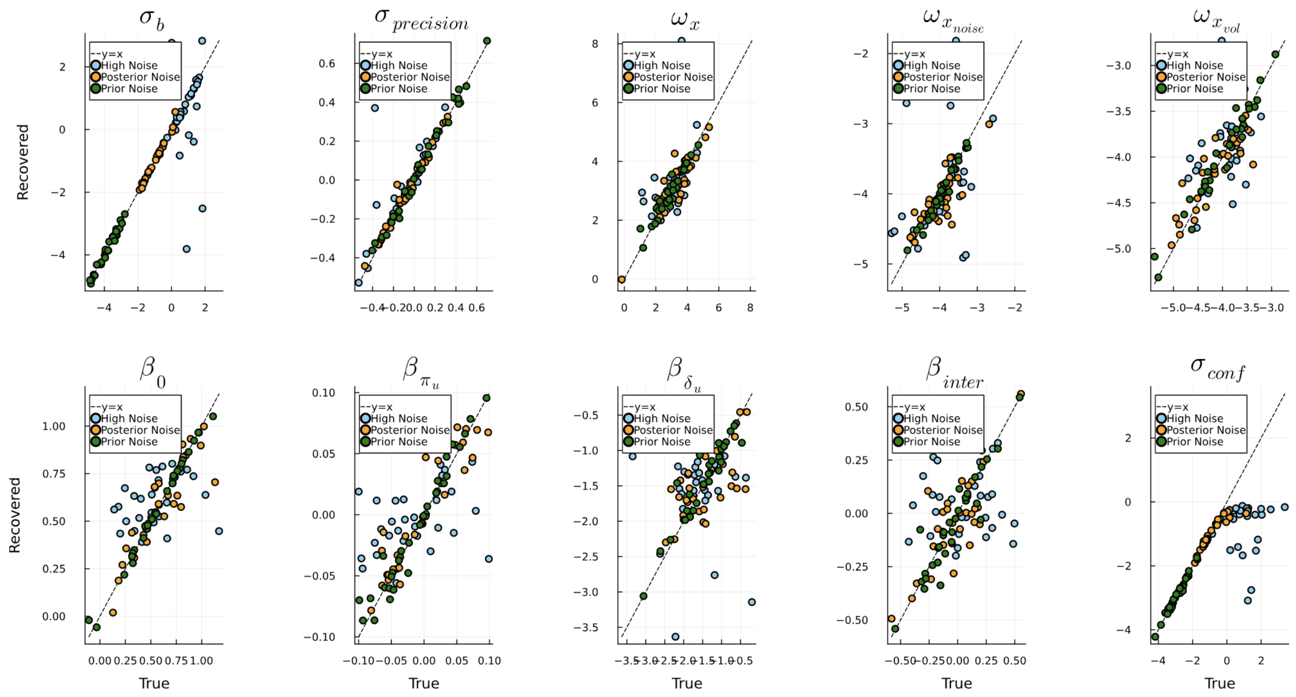


**Figure S3.** Scatter plots of the recovarability of each parameter for the different noise settings.

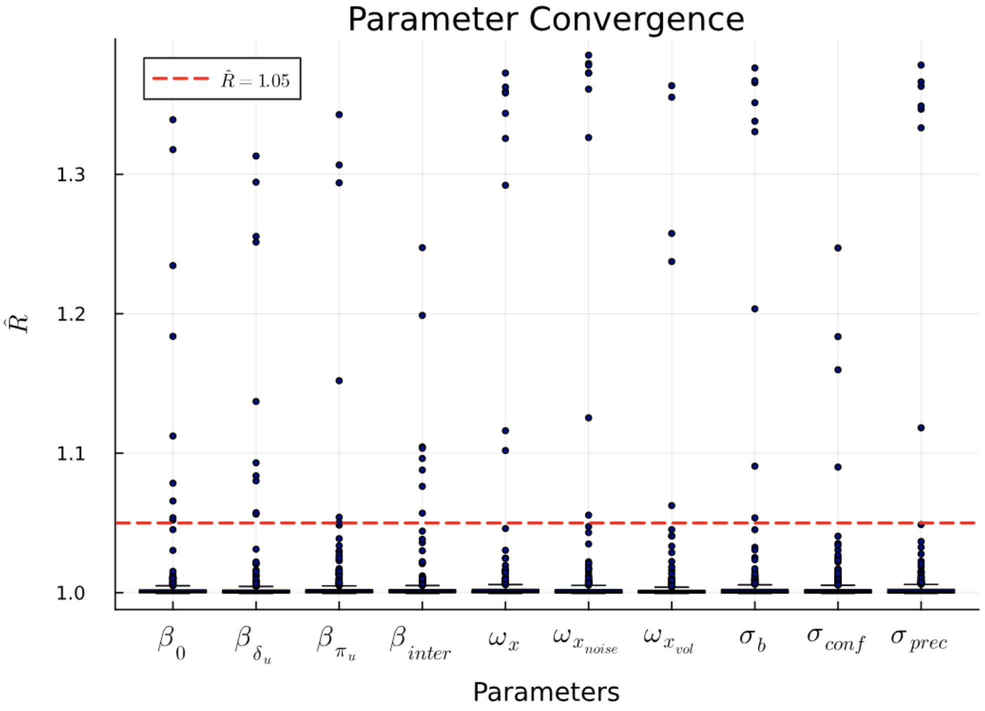


**Figure S4.** Chain convergence analysis using the $\hat{R}$ metric. The plot shows the parameter-averaged $\hat{R}$, the dotted red line represents the $\hat{R} = 1.05$ boundary.

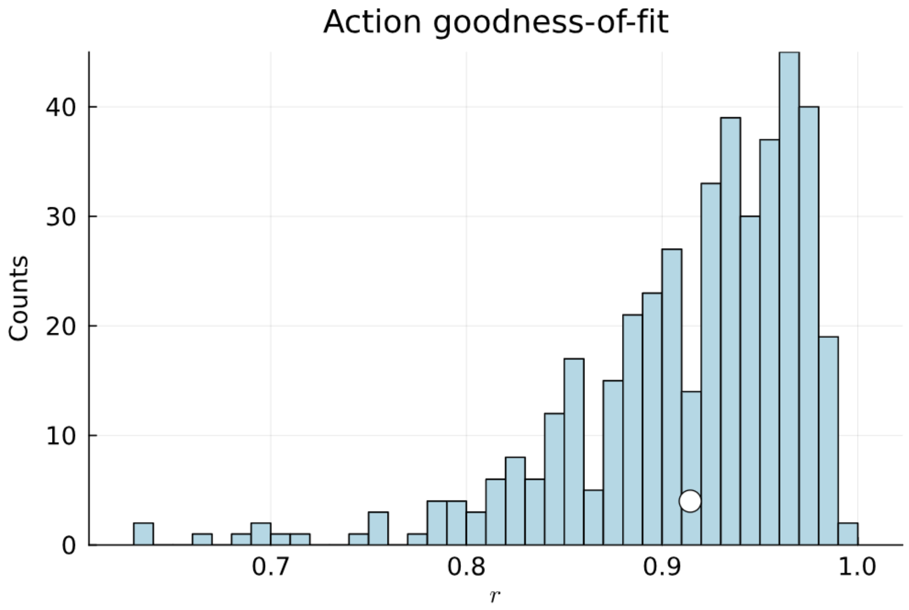


**Figure S5.** Bucket placement (action) goodness-of-fit, as measured by Pearson's correlation coefficient ($r$) between the empirical and the simulated bucket positions.

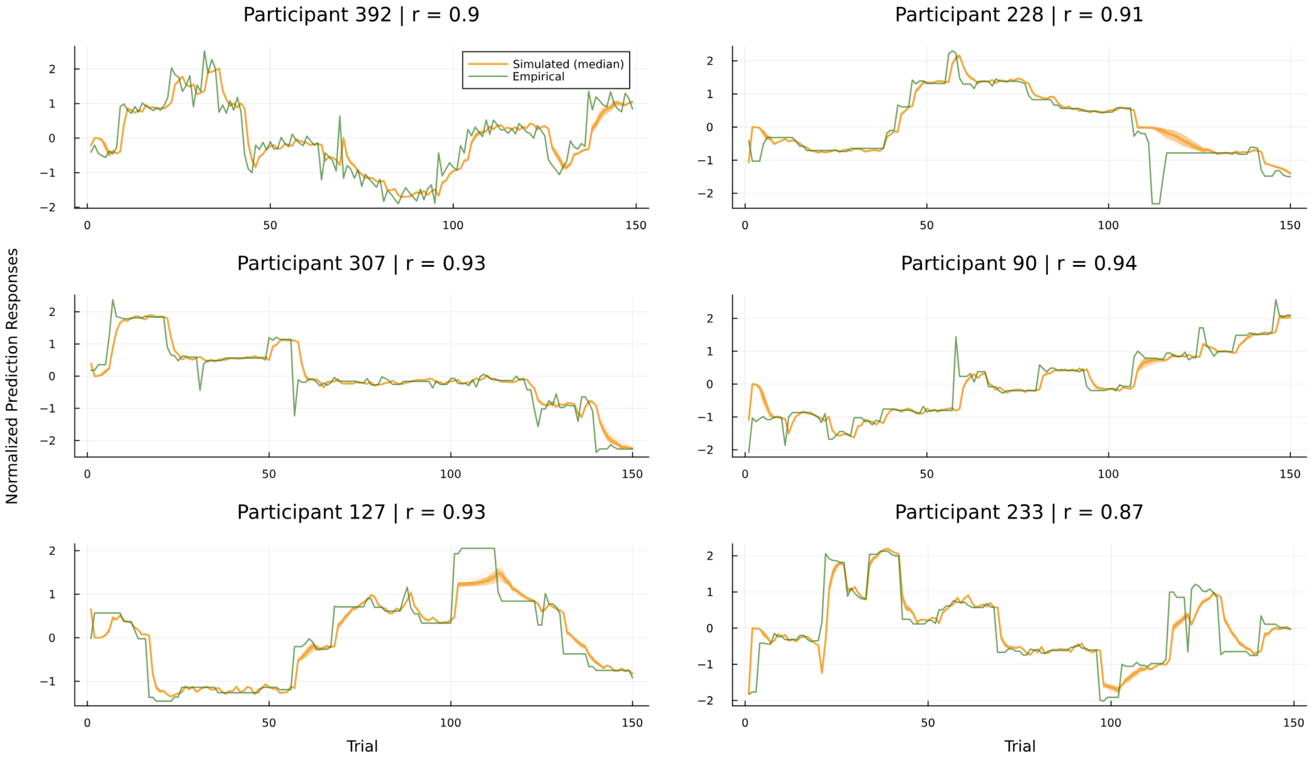


**Figure S6.** Bucket placement (action) model predicted responses. Excellent fits, as measured by Pearson's correlation coefficient ($r$), are obtained. The thick orange line identifies the simulated response from the median of the approximate posterior, while the thinner orange lines show the behaviour generated from the 30 random samples.

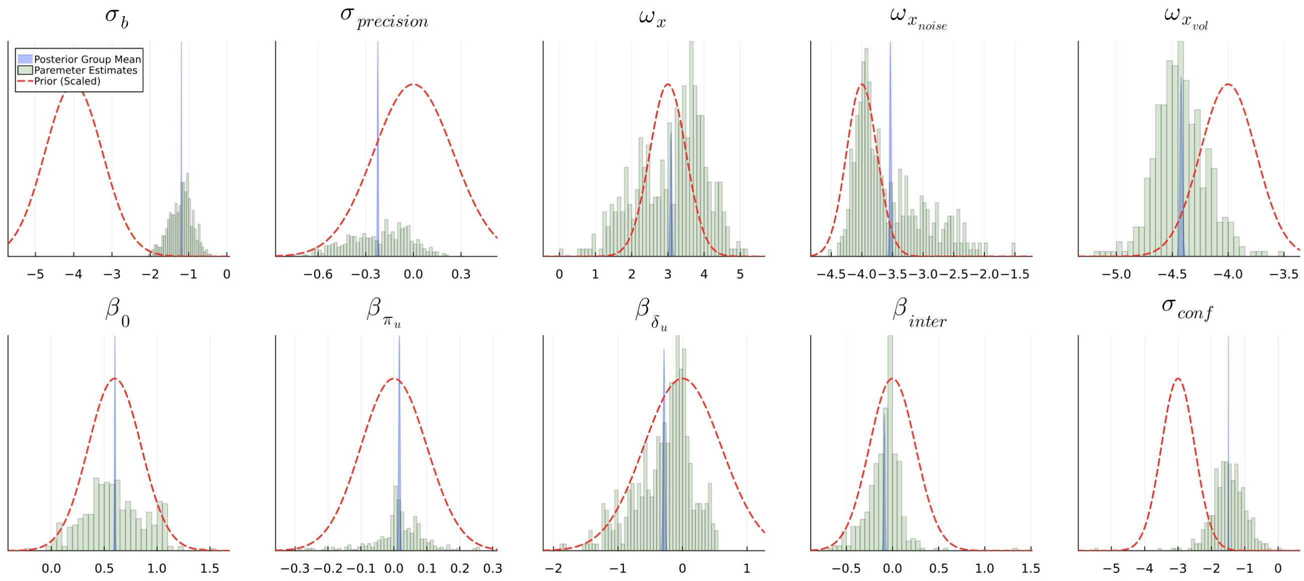


**Figure S7.** Histograms showing the posterior median values for every parameter obtained after model inversion on the full dataset (green), and the posterior group mean (blue). We show on top the initial prior densities (red). Please note that the y axis is not scaled.

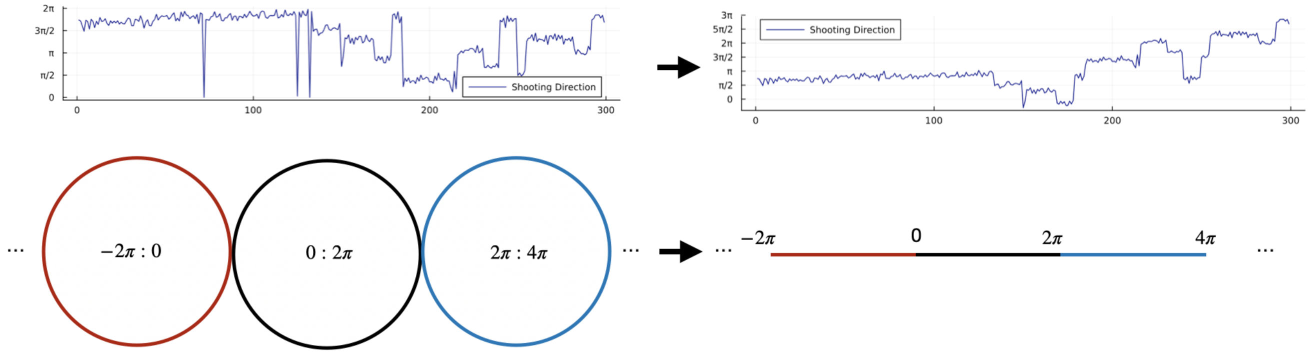


**Figure S8.** Unrolling the circle to enable circular data modelling.

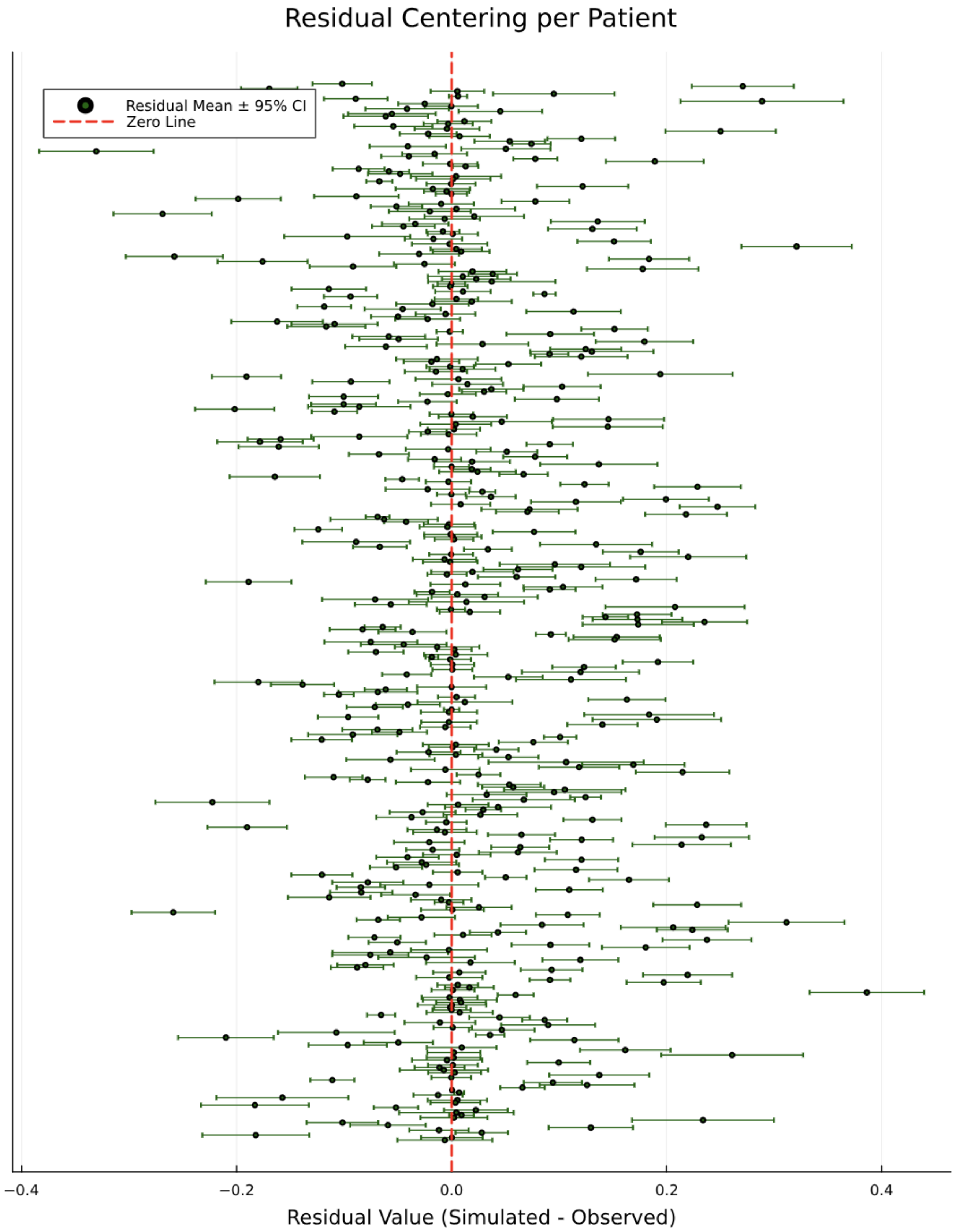


**Figure S9.** Confidence rating fit residuals, single-patient distribution.

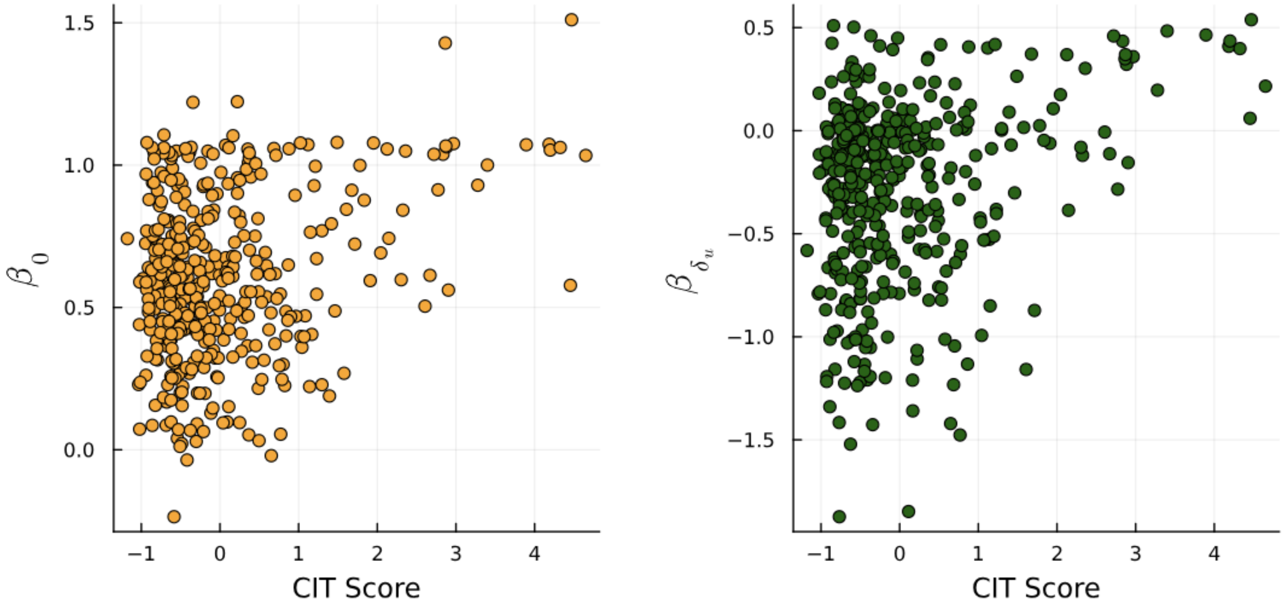


**Figure S10.** Scatter plots of the CIT score against the two significant inverted model parameters: $\beta_0$ and $\beta_{\delta_u}$.
